# Simultaneous CRISPR/Cas9-mediated editing of cassava *eIF4E* isoforms *nCBP-1* and *nCBP-2* reduces cassava brown streak disease symptom severity and incidence

**DOI:** 10.1101/209874

**Authors:** Michael A. Gomez, Z. Daniel Lin, Theodore Moll, Raj Deepika Chauhan, Kelley Renninger, Getu Beyene, Nigel J. Taylor, J. Carrington, B. Staskawicz, R. Bart

## Abstract

<u>C</u>assava <u>b</u>rown <u>s</u>treak <u>d</u>isease (CBSD) is a major constraint on cassava yields in East and Central Africa and threatens production in West Africa. CBSD is caused by two species of positive sense RNA viruses belonging to the family *Potyviridae*, genus *Ipomovirus: <u>C</u>assava <u>b</u>rown <u>s</u>treak <u>v</u>irus* (CBSV) and *<u>U</u>gandan <u>c</u>assava <u>b</u>rown <u>s</u>treak <u>v</u>irus* (UCBSV). Diseases caused by the family *Potyviridae* require the interaction of viral genome-linked protein (VPg) and host <u>e</u>ukaryotic translation <u>i</u>nitiation <u>f</u>actor <u>4E</u> (eIF4E) isoforms. Cassava encodes five eIF4E proteins: eIF4E, eIF(iso)4E-1, eIF(iso)4E-2, <u>n</u>ovel <u>c</u>ap-<u>b</u>inding <u>p</u>rotein-<u>1</u> (nCBP-1), and nCBP-2. Protein-protein interaction experiments consistently found that VPg proteins associate with cassava nCBPs. CRISPR/Cas9-mediated genome editing was employed to generate *ncbp-1, ncbp-2*, and *ncbp-1/ncbp-2* mutants in cassava cultivar 60444. Challenge with CBSV showed that *ncbp-1/ncbp-2* mutants displayed delayed and attenuated CBSD aerial symptoms, as well as reduced severity and incidence of storage root necrosis. Suppressed disease symptoms were correlated with reduced virus titer in storage roots relative to wild-type controls. Our results demonstrate the ability to modify multiple genes simultaneously in cassava to achieve tolerance to CBSD. Future studies will investigate the contribution of remaining eIF4E isoforms on CBSD and translate this knowledge into an optimized strategy for protecting cassava from disease.

## Introduction

<u>C</u>assava <u>b</u>rown <u>s</u>treak <u>d</u>isease (CBSD) is a threat to food and economic security for smallholder farmers in sub-Saharan Africa. First reported in the 1930s in lowland and coastal East Africa, CBSD has since spread west to higher altitudes in Uganda, Kenya, Tanzania, Burundi, and the Democratic Republic of Congo (Adams *et al.*, 2013; Alicai *et al.*, 2007; Bigirimana *et al.*, 2011; Mbanzibwa *et al.*, 2011; Mulimbi *et al.*, 2012). The CBSD vector is the whitefly *Bemisia tabaci* which has a broad geographical distribution across sub-Saharan Africa (Legg *et al.*, 2014). CBSD symptoms include leaf chlorosis, brown streaks on stems, and necrosis of the storage roots. CBSD immunity, or complete non-infection of the cassava plant *(Manihot esculenta* Crantz), has not been observed within known farmer cultivars (Kaweesi *et al.*, 2014). Infection can occur in resistant cultivars such as Kaleso and Namikonga, but multiplication, movement, and disease symptoms are limited (Kaweesi *et al.*, 2014). Tolerant cultivars Nachinyaya and Kiroba can be infected and support virus movement and replication, but with intermediate symptoms, while susceptible cassava cultivars 60444 and Albert support high levels of virus concentration and develop severe CBSD symptoms (Hillocks *et al.*, 2001; Maruthi *et al.*, 2014; Masiga *et al.*, 2014; Ogwok *et al.*, 2015). Since symptoms may be subtle or develop only within the underground storage roots, CBSD may claim an entire crop without the farmer’s knowledge until harvest (Legg *et al.*, 2015; Patil *et al.*, 2015). Necrotic lesions render the storage roots unfit for market and human consumption with losses of up to 70% root weight reported (Hillocks *et al.*, 2001). The international institute of Tropical Agriculture (IITA) estimated that CBSD causes $175 million loss in East Africa each year (Michael, 2013).

The causative agents of CBSD, *<u>C</u>assava <u>b</u>rown <u>s</u>treak <u>v</u>irus* (CBSV) and *<u>U</u>gandan <u>c</u>assava <u>b</u>rown <u>s</u>treak <u>v</u>irus* (UCBSV), belong to the family *Potyviridae* (Genus: *Ipomovirus)* (Revers and García, 2015). These non-enveloped, flexuous, filamentous viruses contain a positive-sense, single-stranded RNA, with a 3’-poly(A) terminus (King *et al.*, 2012). The CBSV genome encodes a polyprotein of 2902 amino acids that is proteolytically cleaved into 10 mature proteins (Mbanzibwa *et al.*, 2009). A viral genome-linked (VPg) protein is covalently linked to the 5’ end of the viral genome and is required for infection by this pathogen (Robaglia and Caranta, 2006; Wang and Krishnaswamy, 2012).

Resistance to plant pathogens can be controlled either through dominant or recessive gene inheritance. Resistance genes encoding nucleotide-binding leucine-rich repeat receptors, which are dominant sources of extreme resistance against adapted pathogens in many pathosystems, have been cloned and characterized for potyviral diseases, but an overrepresentation in recessive resistance to potyviruses is well documented (de Ronde et al., 2014; Revers and Nicaise, 2014). Recessive resistance to potyviruses is typically associated with mutations in the <u>e</u>ukaryotic translation initiation factor <u>4E</u> (eIF4E) protein family (Bastet *et al.*, 2017; Robaglia and Caranta, 2006). Ethyl methanesulfonate- and transposon-mutagenesis screens in *Arabidopsis thaliana* for decreased susceptibility to <u>Tu</u>rnip <u>M</u>osaic poty<u>v</u>irus (TuMV) identified *eIF(iso)4E* as a loss of susceptibility locus (Duprat *et al.*, 2002; Lellis *et al.*, 2002). More broadly, polymorphisms in eIF4E isoforms of pepper, tomato, lettuce, pea, and other crops confer resistance to numerous potyviruses (Robaglia and Caranta, 2006). The direct physical interaction between potyvirus VPg and specific host eIF4E isoforms is well supported through *in vitro* and *in vivo* binding assays (Kang *et al.*, 2005; Leonard *et al.*, 2000; Schaad *et al.*, 2000; Wittmann *et al.*, 1997; Yeam *et al.*, 2007). In most of these cases, amino acid substitutions within the interaction domains on either VPg or eIF4E isoforms abolished infection, highlighting the necessity of eIF4E isoform interaction.

The eIF4E protein family plays an essential role in the initiation of cap-dependent mRNA translation. eIF4E isoforms interact with the 5’-7mGpppN-cap of mRNA and subsequently recruit a complex of initiation factors for ribosomal translation. eIF4E and its different isoforms, eIF(iso)4E and novel cap-binding protein (nCBP), vary in degrees of functional redundancy and may have undergone neo- or subfunctionalization (Browning and Bailey-Serres, 2015). Little is known regarding nCBPs, in particular. Studies in *A. thaliana* have shown that nCBP exhibits weak cap-binding, similar to eIF(iso)4E, and increased levels in cap-binding complexes at early stages of cell growth (Kropiwnicka *et al.*, 2015, Bush *et al.*, 2009). However, the specialized function of plant nCBPs remains unknown. Potyviruses hijack the eIF4E protein family via their VPg for translation initiation, genome stability, and/or viral movement (Fig S1) (Contreras-Paredes *et al.*, 2013; Eskelin *et al.*, 2011; Gao *et al.*, 2004; Miras *et al.*, 2017; Zhang *et al.*, 2006). Transgenic approaches leveraging amino acid changes that abolish interaction with VPg or loss of the VPg-associated eIF4E protein have previously been implemented as a form of potyviral disease control (Cui and Wang, 2017; Piron *et al.*, 2010; Wang, 2015).

Targeted genome editing techniques have emerged as alternatives to classical plant breeding and transgenic methods (Belhaj *et al.*, 2015). The CRISPR (<u>C</u>lustered <u>R</u>egularly <u>i</u>nterspaced <u>S</u>hort <u>P</u>alindromic <u>R</u>epeats)/Cas9 (<u>C</u>RISPR <u>a</u>ssociated <u>p</u>rotein <u>9</u>) system has rapidly become a favored tool for biotechnology because of its simple design and easy construction of reagents. The Cas9 nuclease is recruited to a specific site within the genome via a guide RNA (gRNA) (Jinek *et al.*, 2012). Upon binding, Cas9 induces a <u>d</u>ouble <u>s</u>trand <u>b</u>reak (DSB) at the target site (Belhaj *et al.*, 2015). Repair of the DSB by error-prone <u>n</u>on-<u>h</u>omologous <u>e</u>nd joining (NHEJ) can generate <u>in</u>sertion or <u>del</u>etion (INDEL) mutations that disrupt gene function by altering the reading frame and/or generate a premature stop codon (Britt, 1999; Gorbunova and Levy, 1999). We aimed to apply the CRISPR/Cas9 technology to knockout the VPg-associated cassava eIF4E isoform(s). This approach to engineering potyvirus resistance has been successfully demonstrated in *A. thaliana* and cucumber (Chandrasekaran *et al.*, 2016; Pyott *et al.*, 2016). Here, we show that targeted mutagenesis of specific cassava *eIF4E* isoforms *nCBP-1* and *nCBP-2* by the CRISPR/Cas9 system reduces levels of CBSD associated disease symptoms and CBSV accumulation in storage roots. Simultaneous disruption of both *nCBP* isoforms resulted in a larger decrease in disease symptoms than disruption of either isoform individually.

## Results

### Identification and sequence comparison of eIF4E isoforms in cassava varieties

To identify the eIF4E family protein(s), a BLAST search of the AM560-2 cassava cultivar genome (assembly version 6.1) was done via Phytozome using *A. thaliana* eIF4E family proteins as the queries (Bredeson *et al.*, 2016; Goodstein *et al.*, 2012). Five cassava proteins were found that phylogenetically branched with the eIF4E, eIF(iso)4E, and nCBP sub-groups (Fig. 1a). Two of the cassava eIF4E family proteins joined within the eIF(iso)4E sub-group, and another two joined within the nCBP sub-group. This is in agreement with findings by Shi *et al.* (2017). Percent identity analysis further supported this grouping as the eIF(iso)4E- and nCBP-similar proteins had high amino acid identity (Fig. 1b). Based upon this phylogenetic analysis, one *eIF4E*, two *eIF(iso)4E*, and two *nCBP* genes cassava genes were re-named according to their sub-groups (Fig. 1c).

**Figure 1.**
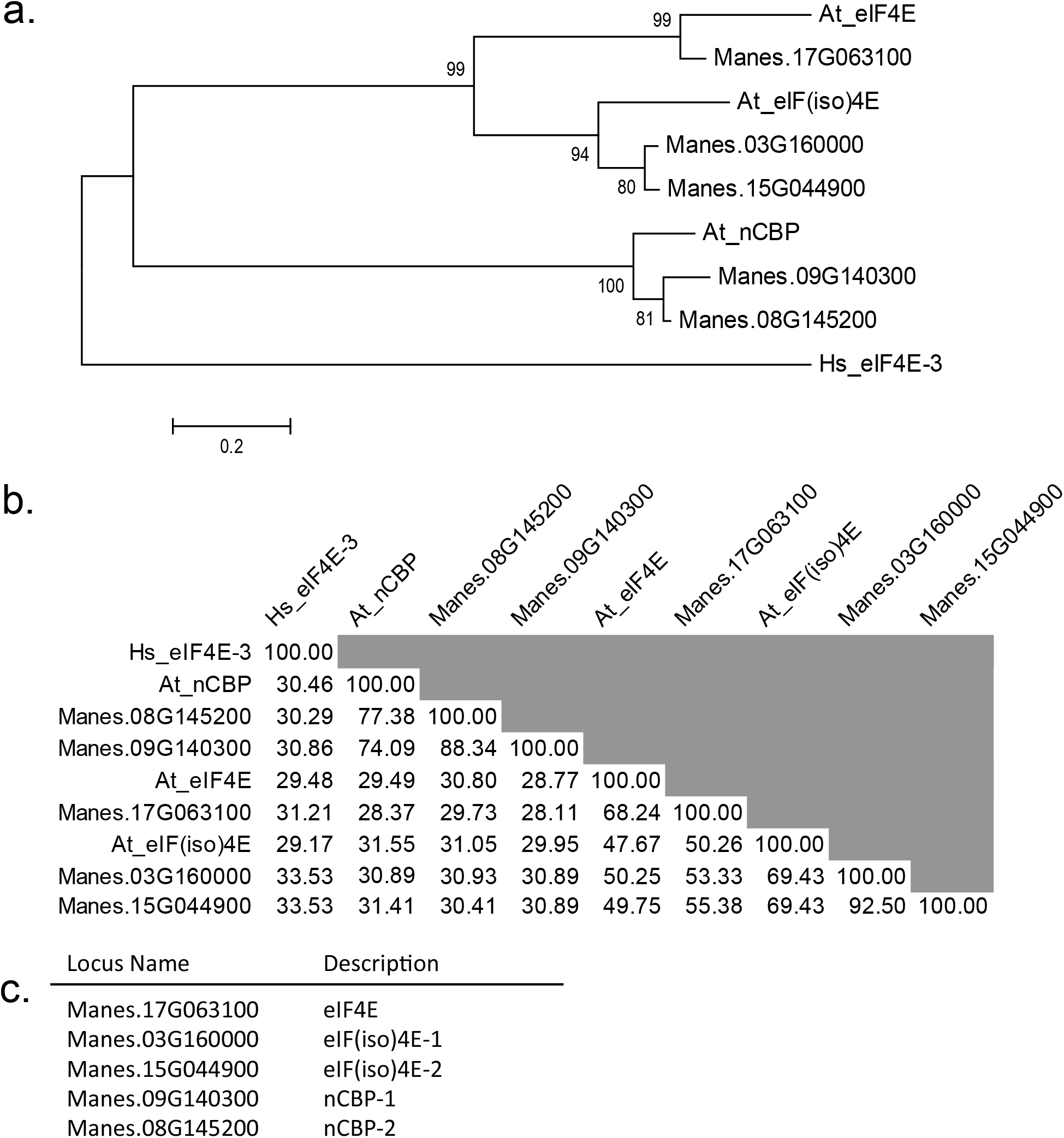
Identification of cassava eIF4E family homologs. (a) Phylogenetic relationships of cassava eIF4E family with Arabidopsis thaliana, At, eIF4E family inferred by the Maximum Likelihood method (Le and Gascuel, 1993). The percentage of trees in which the associated taxa clustered together is shown next to the branches. The tree is drawn to scale, with branch lengths measured in the number of substitutions per site. Tree is rooted to Homo sapiens, Hs, eIF4E-3. (b) Percent identity matrix of cassava eIF4E family with At eIF4E family using amino acid sequences. (c) Descriptions of cassava eIF4E family based upon phylogenetic relationships and percent identity matrix.

### Multiple cassava eIF4E isoforms interact with VPg

We investigated CBSV VPg association with cassava eIF4E isoforms through co-immunoprecipitation (co-IP) experiments. Several attempts to co-express and co-IP tagged forms of CBSV VPg and cassava eIF4E isoforms in *Nicotiana benthamiana* were unsuccessful, due possibly to very low levels of VPg expression or competitive interference from endogenous eIF4E isoforms (Avila et al., 2015, Monger et al., 2001). As an alternative, tests for interaction were done using bacterially expressed 6xHIS-VPg-6xHIS-3xFLAG protein mixed with lysates of *Nicotiana benthamiana* expressing YFP-eIF4E isoforms. This approach was first validated using the well characterized TuMV VPg association with *Arabidopsis* eIF(iso)4E (Fig. S2). Interactions were tested between tagged forms of CBSV-Naliendele isolate TZ:Nal3-1:07 (CBSV-Nal) VPg and cassava eIF4E isoforms. The results indicate that CBSV-Nal VPg can associate with all YFP-fused cassava eIF4E isoforms but not YFP alone (Fig. 2a).

**Figure 2.**
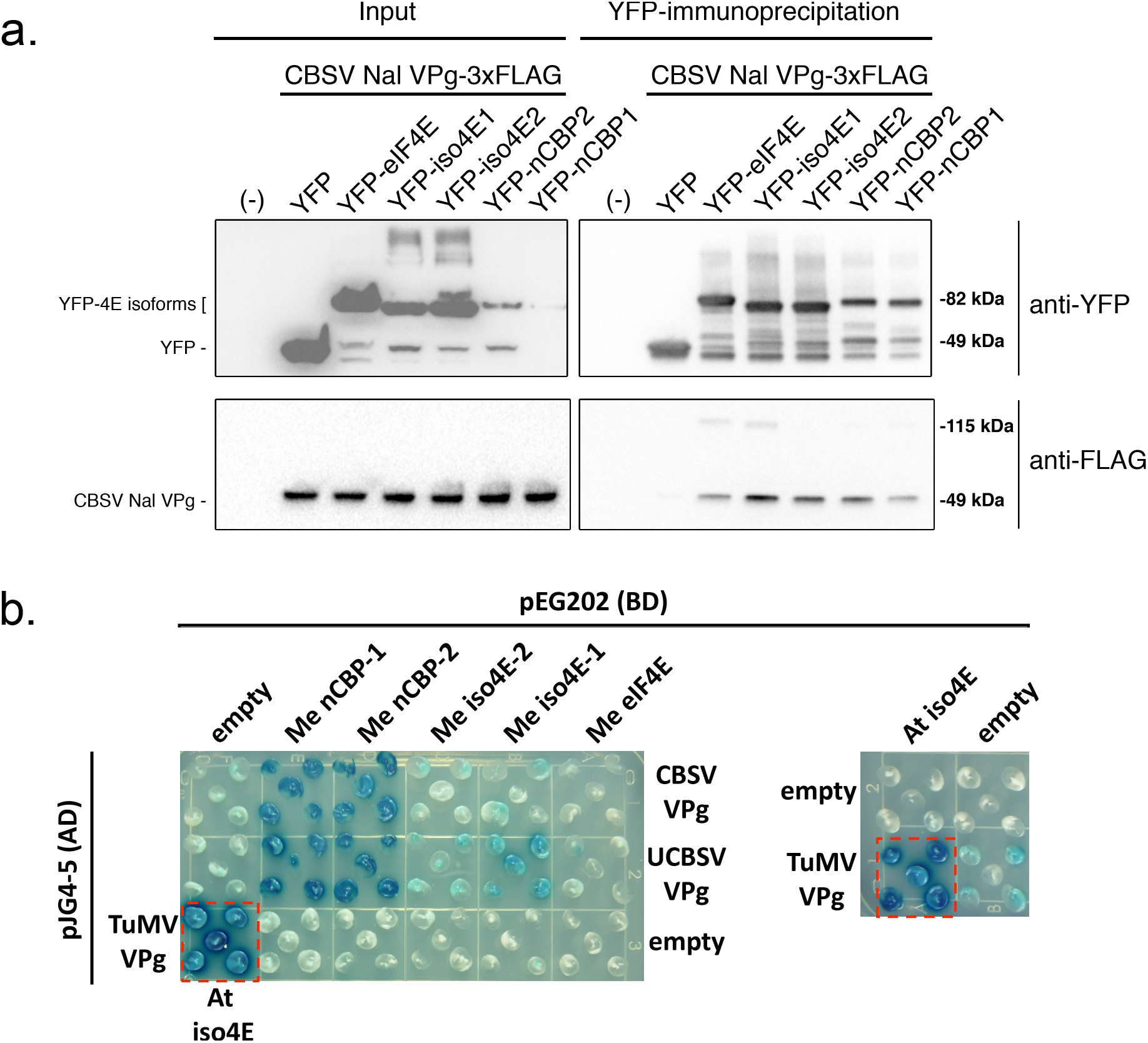
CBSV and UCBSV VPg’s interact with cassava nCBP-1 and nCBP-2. (a) Immunoprecipitation of YFP-cassava eIF4E isoform fusions, expressed in Nicotiana benthamiana, co-immunoprecipitate purified CBSV-Nal VPg-3xFLAG that was added to plant extracts. (b) Yeast two-hybrid constructs consist of B42 activation domain (AD) fused to the CBSV Naliendele VPg and UCBSV T04 VPg, and LexA DNA binding domain (BD) bound to cassava eIF4E family members. Blue coloration represents β-galactosidase activity from activation of lacZ reporter gene by protein-protein interaction. Five yeast transformants are displayed on the dropout medium SD Gal/Raf SD-UTH. Positive control is shown in the dashed red box (TuMV VPg-AD and Arabidopsis thaliana eIF(iso)4E-BD).

As a complementary approach to the described co-IP experiments, a yeast two-hybrid system was used to assess the VPg-eIF4E isoform interactions. The VPg proteins from CBSV-Nal and UCBSV isolate UG:T04-42:04 (UCBSV-T04) were fused to the B42 activation domain and transformed into yeast strain EGY48. All five cap-binding proteins were fused to the LexA DNA-binding domain and transformed into VPg yeast lines. Likewise fused, TuMV VPg and *A. thaliana* eIF(iso)4E were transformed into yeast as a positive control, and empty vectors were transformed as negative controls. Five colonies from each transformation were plated on selective media supplemented with X-gal. Protein-protein interaction dependent activity of the β-galactosidase reporter was indicated by a blue color. Both nCBP-1 and nCBP-2 showed strong interactions with the VPgs based on color intensity, and comparable to the positive control (Fig. 2b). The eIF(iso)4E proteins appeared to exhibit a weak interaction with VPg proteins, while eIF4E-VPg interactions were not detected (Fig. 2b).

### Site-specific mutation of *nCBP* isoforms by transgenic expression of sgRNA-guided Cas9

Taken together, the protein-protein interaction data suggest that viral VPg proteins can interact with multiple members of the cassava eIF4E family. The nCBP clade consistently interacted with CBSV VPg and was prioritized for functional characterization. CRISPR/Cas9 was employed to generate mutant alleles of cassava *nCBP* isoforms. Five constructs were assembled to target various sites in *nCBP-1, nCBP-2*, and both genes simultaneously (Table 1). *Agrobacterium* carrying these constructs were then used to transform <u>f</u>riable <u>e</u>mbryogenic <u>c</u>alli (FEC) derived from cassava cultivar 60444 (Fig. S3). Multiple independent T0 transgenic plant lines were recovered for each construct (Table S1). Sites in each *nCBP* gene were targeted to disrupt restriction enzyme recognition sequences (Fig. S4). Restriction digestion analysis of PCR products from T0 plants with SmlI detected mutagenesis of *nCBP* genes (Fig. S4).

**Table 1.**
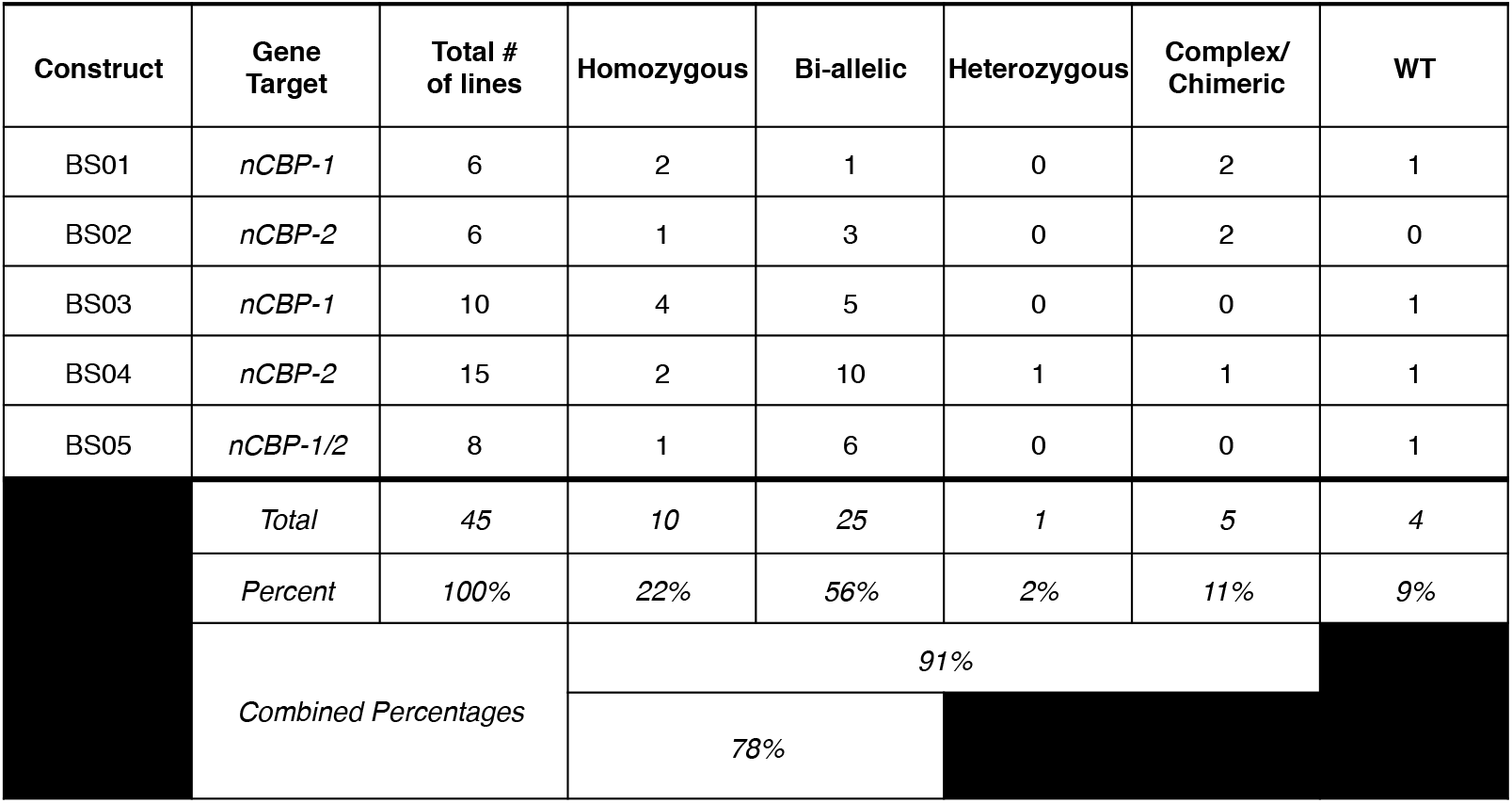
Genotype counts of transgenic T_0_ cassava lines.

| Construct | Gene Target | Total # of lines | Homozygous | Bi-allelic | Heterozygous | Complex/ Chimeric | WT |
| --- | --- | --- | --- | --- | --- | --- | --- |
| BS01 | *nCBP-1* | 6 | 2 | 1 | 0 | 2 | 1 |
| BS02 | *nCBP-2* | 6 | 1 | 3 | 0 | 2 | 0 |
| BS03 | *nCBP-1* | 10 | 4 | 5 | 0 | 0 | 1 |
| BS04 | *nCBP-2* | 15 | 2 | 10 | 1 | 1 | 1 |
| BS05 | *nCBP-1/2* | 8 | 1 | 6 | 0 | 0 | 1 |
|  | *Total* | 45 | 10 | 25 | 1 | 5 | 4 |
|  | *Percent* | 100% | 22% | 56% | 2% | 11% | 9% |
|  | *Combined Percentages* |  | 91% |  |  |  |  |
|  |  |  | 78% |  |  |  |  |

The range of mutations generated in each transgenic plant was analyzed by subcloning and sequence analysis, revealing an array of homozygous, bi-allelic, heterozygous, complex, and wild-type genotypes (Table S1). Bi-allelic mutations contained different mutations on the two alleles. Heterozygous plants carried one mutagenized allele and one wild-type allele. Plants were considered complex if they carried more than two sequence patterns, strongly suggesting chimerism (Odipio *et al.*, 2017; Zhang *et al.*, 2014). The genotypes of edited plants had Cas9-induced INDELs ranging from insertions of 1 to 16 bp and deletions as large as 127 bp (Table S1). Review of all genotyped plants revealed that 10/45 (22%) carried homozygous mutations, 25/45 (56%) carried bi-allelic mutations, 1/45 (2%) were heterozygous, 5/45 (11%) were complex, and another 4/45 (9%) were wild-type genotypes (Table 1). In total, 78% of plants contained either homozygous or bi-allelic mutations, and CRISPR/Cas9 activity was observed in 91% of the plants studied.

### Sequence analysis of INDEL-induced frameshifts in *nCBP*s identifies multiple *ncbp-1* splice variants

Edited lines with mutations in *nCBP-1* and *nCBP-2* individually, as well as both *nCBPs* in tandem, were selected for CBSV disease trials in a greenhouse. Lines with homozygous mutations in exon 1 were prioritized (Table S1). The mutant lines chosen for these trials, *ncbp-1* #1, *ncbp-2* #6, *ncbp-1/2* #2, and *ncbp-1/2* #8, each had an INDEL at the 3’ end of the first exon of each targeted gene (Fig. 3). The INDELs either directly resulted in a frameshift or disrupted the exon-intron splice sites so that an out-of-frame splice variant was predicted to be produced (Fig. 4A, S5). Predicted out-of-frame splice variants from homozygous mutants were validated by sequencing of cDNA amplicons. To characterize bi-allelic mutations, cDNA clone sequencing (clone-seq) was done (Fig. 4B). The homozygous *ncbp-1* allele from *ncbp-1/2* #2 was also analyzed for comparison. Of nine *ncbp-1* cDNA clones from mutant line *ncbp-1/2* #2, eight displayed the wild-type splicing pattern (referred to as type 1) with the insertion of an A from genomic DNA sequence results. This generates a frameshift, culminating in a premature stop codon. An alternative splice variant (referred to as type 2) was also observed (Fig. 4B). This variant results in retention of 35 nucleotides from intron 1 but does not shift the reading frame. Thus, this splice variant encodes a full protein with a 12 amino acid internal insertion. This splicing pattern was not observed in any wild-type *nCBP-1* clones, however, may occur at low frequency.

**Figure 3.**
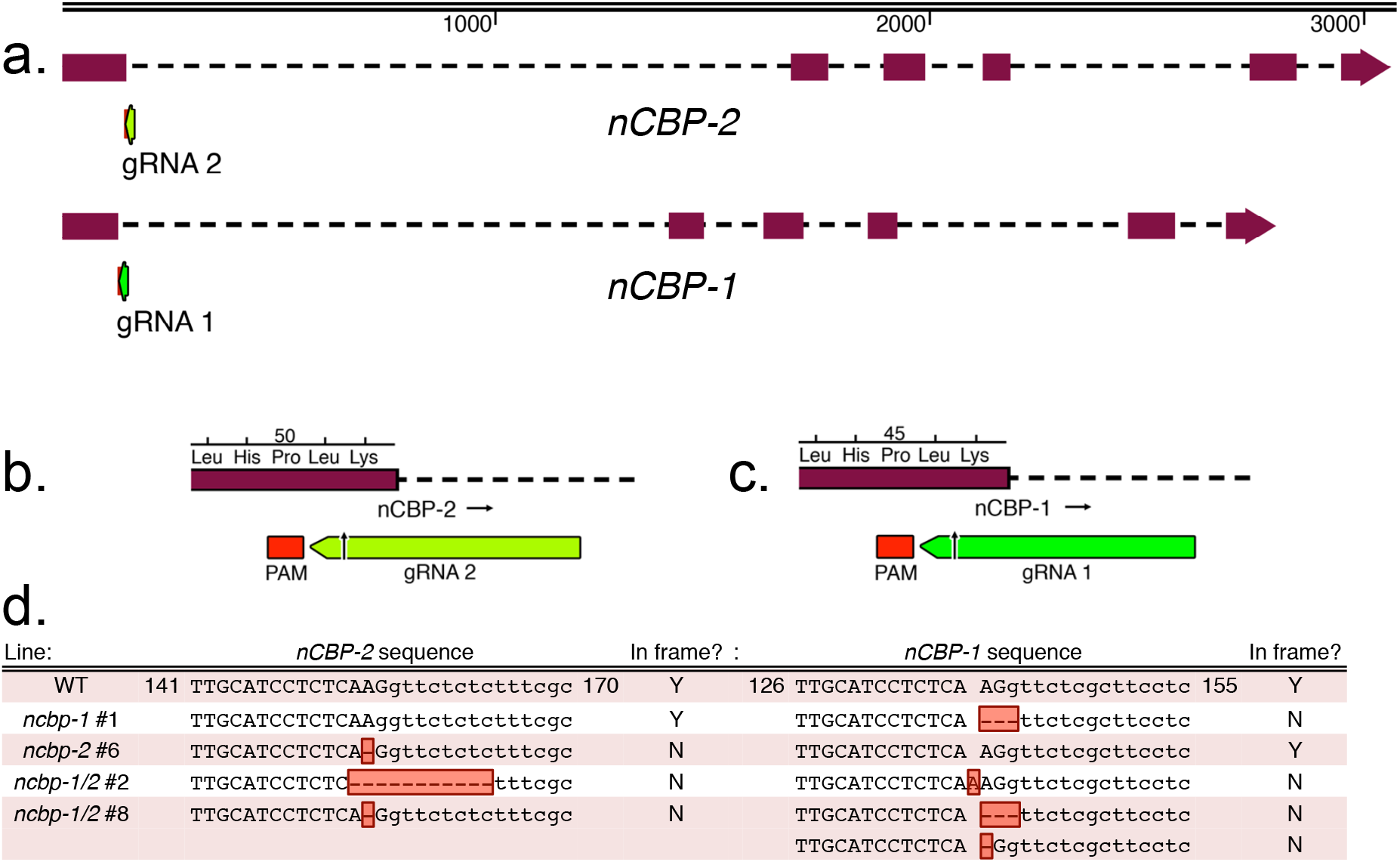
Cas9 induces INDELs at *nCBP-1* and *nCBP-2* gRNA target sites in BS05 transgenic lines. (a) Sequences at the junction of the first exon-intron boundary were selected for targeting of the Cas9 nuclease. Lengths of *nCBP-1* and *nCBP-2* genes are to nucleotide scale (top bar). Exons are denoted by solid blocks and introns are represented as dashed lines. Arrowheads indicate the 3’ terminus. (b) Diagram of the protospacer adjacent motif (PAM) and guide RNA (gRNA) targeting *nCBP-2.* (c) Diagram of the PAM and gRNA targeting *nCBP-1.* (d) INDELs of *nCBP-1* and *nCBP-2* in mutant lines BS01 #1, BS02 #6, BS05 #2, and BS05 #8. Upper and lower cases denote exonic and intronic sequence, respectively. Red boxes indicate INDELs.

**Figure 4.**
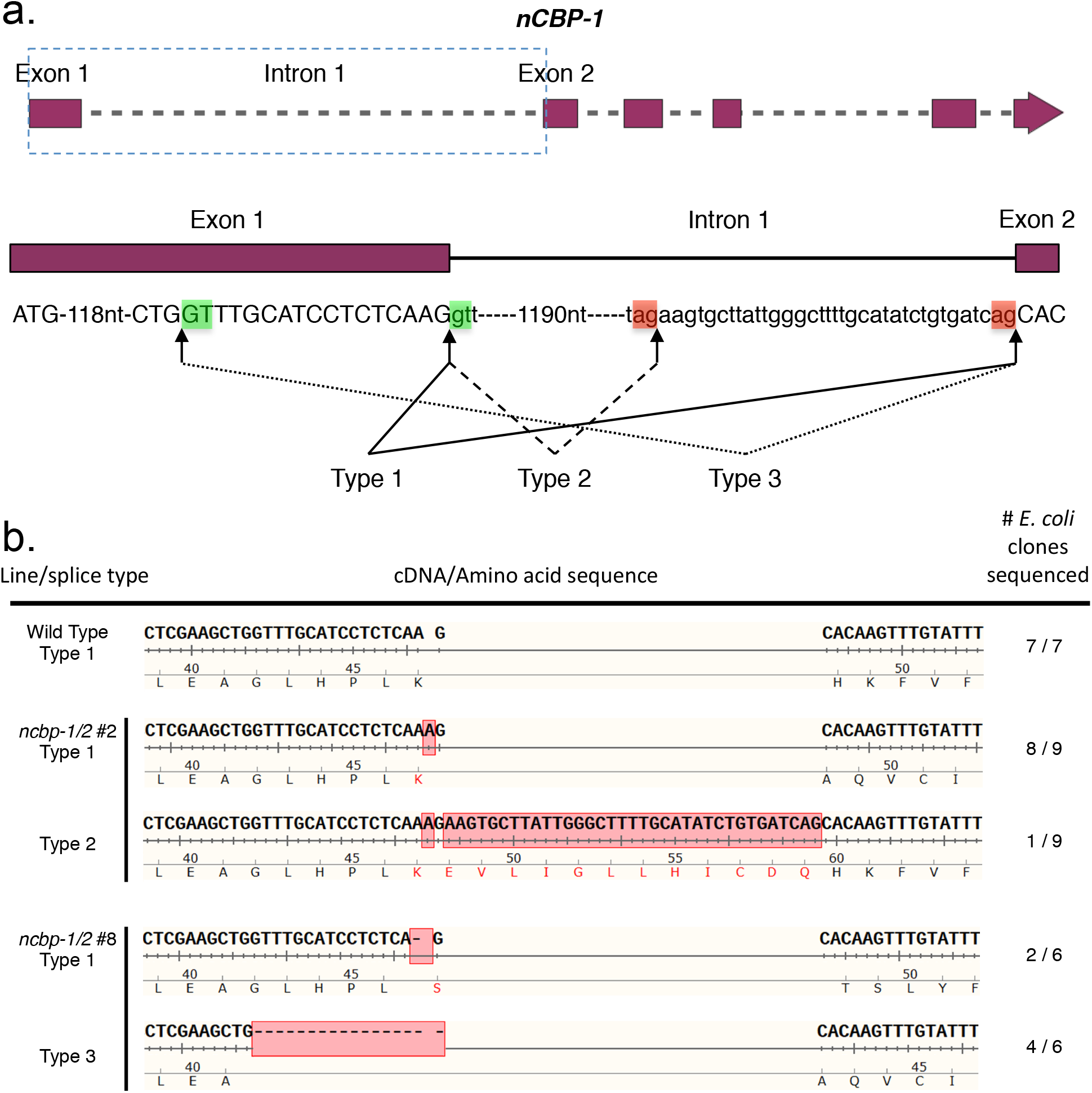
Alternative splicing of *ncbp-1* alleles is detected in *ncbp-1/2* double mutants. (a) Schematic of canonical and alternative *nCBP-1* splice sites. Boxed region of the *nCBP-1* gene model is enlarged below. Exon and intron sequences are given in capital and small letters, respectively. Green and red boxes highlight splice motifs at the 5’ and 3’ end of introns, respectively. Type 1 splicing produces the predicted wild type *nCBP-1* cDNA sequence. Type 2 and 3 splicing are observed in ncbp-1/2 lines #2 & #8, respectively. (b) cDNA sequences detected in clone-seq experiments. Red boxes denote INDELs resulting from both CRISPR/Cas9-mediated edits and alternative splicing. In *ncbp-1/2* #2, type 2 splicing results in retention of 3’ sequence from intron 1 of ncbp-1 (1 of 9 clones sequenced). In *ncbp-1/2* #8, an INDEL disrupting the canonical splice motif between exon 1 and intron 1 of *ncbp-1* results in a type 3 splice variant (4 of 6 clones sequenced).

Clone-seq analysis of *ncbp-1* transcripts from mutant line *ncbp-1/2* #8 cDNA similarly found predicted INDELs. Two cDNA clones displayed the wild-type (type 1) splicing pattern and the predicted deletion of an A. Four clones showed a sequence pattern that suggests a third splicing variant (type 3) at an upstream alternative splice site (Fig. 4) (Reddy *et al.*, 2007). Both observed cDNA sequence patterns are frameshifted.

### Off-target analysis

A full genome assembly for cassava variety 60444 is not available. However, Illumina resequencing data for this variety was recently published (Bredeson et al., 2016). Using these data we created an AM560-2 v6 reference-based assembly for 60444 and used it to predict potential off-targets for gRNA1 (target *nCBP-1)* and gRNA2 (target *nCBP-2)* with the CasOT tool (Xiao et al., 2014; Table S2, Table S3). The top five hits with the fewest mismatches for each gRNA were analyzed with PCR followed by Sanger sequencing. gRNA1 and gRNA2 share several off-targets due to homology between targeted regions, resulting in a combined set of eight off-targets (A-H). Sequence analysis revealed wildtype sequence for off-targets A, B, C, E, F, and H. We were unable to amplify off-target D. We observed a mutation within one allele of G, an off-target for gRNA2. This off-target does not fall within an annotated gene in the cassava version 6 genome. However, gene expression data available through Phytozome suggests this site may be within an intron. The mutated allele had two mismatched nucleotides from gRNA2, while the wild-type allele had an additional third mismatch.

### ncbp-1/ncbp-2 double mutants exhibited reduced CBSD symptom incidence and severity in aerial tissues

The ncbp-1, ncbp-2, ncbp-1/2 #2, ncbp-1/2 #8, and wild-type 60444 plants were chip-bud graft inoculated with CBSV-Nal and monitored for differences in symptom development (Wagaba et. al, 2013). Aerial disease incidence was scored every week for 12 to 14 weeks after grafting and the percentage of plants showing any level of foliar or stem symptoms was recorded at each time point. Fluctuations in the percentage of plants that exhibited symptoms at each time point (% incidence) result from the shedding of symptomatic leaves throughout the experiment. At least five replicate plant clones were included for each genotype (n≥5). Across all three experimental replicates, ncbp-1/ncbp-2 double mutants exhibited delayed symptom development relative to wild type and ncbp-1 (Fig. 5a, S6). ncbp-2 exhibited symptom incidence development similar to wild type and ncbp-1 in two experiments and displayed an intermediate phenotype in the third experiment (Fig 5b, S6).

**Figure 5.**
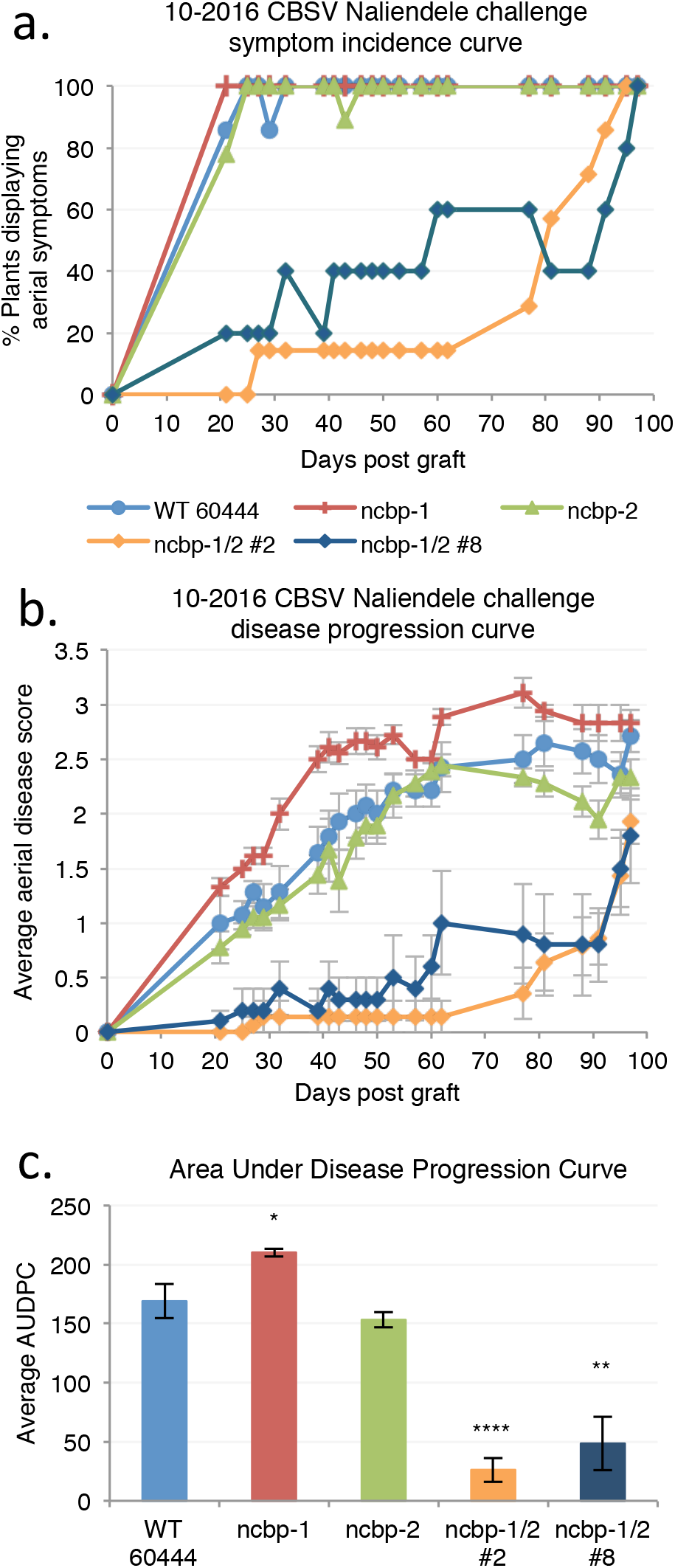
*ncbp-1 ncbp-2* double mutants exhibit delayed CBSV symptom onset and reduced symptom severity. (a), aerial symptom incidence reported as percent of wild type, ncbp-1, ncbp-2, or *ncbp-1 ncbp-2* plants bud-graft inoculated with CBSV Naliendele isolate. *ncbp-1 ncbp-2* double mutant lines #2 and #8 are the product of independent transgenic events. (b), disease progression curves for previously described CBSV inoculated plants. Leaf and stem symptoms were each scored on a 0-4 scale and an average aerial score is shown. (c), average area under the disease progression curve (AUDPC) derived from data plotted in (c). Error bars in (c) and (d) indicate standard error of the mean. Statistical differences were detected by Welch’s t-test, n≤5, α=0.05, *≤0.05, **≤0.01, ****≤0.0001.

In addition to disease incidence, aerial tissue symptom severity was assessed for the CBSV challenges (Table S4, Fig. 5b, 6, S7). Compared to wild-type, the ncbp-1/ncbp-2 double mutants had greatly reduced CBSD severity in all three trials. Area under the disease progression curve (AUDPC) analysis revealed this to be a statistically significant difference in all three experimental replicates (Fig. 5c, S7). Despite the robust stem phenotypes in these experiments, differences in leaf symptoms was variable across experiments (Fig. S8, S9). qPCR analysis of leaf virus titer at the end of challenges also proved highly variable (Fig.S10).

**Figure 6.**
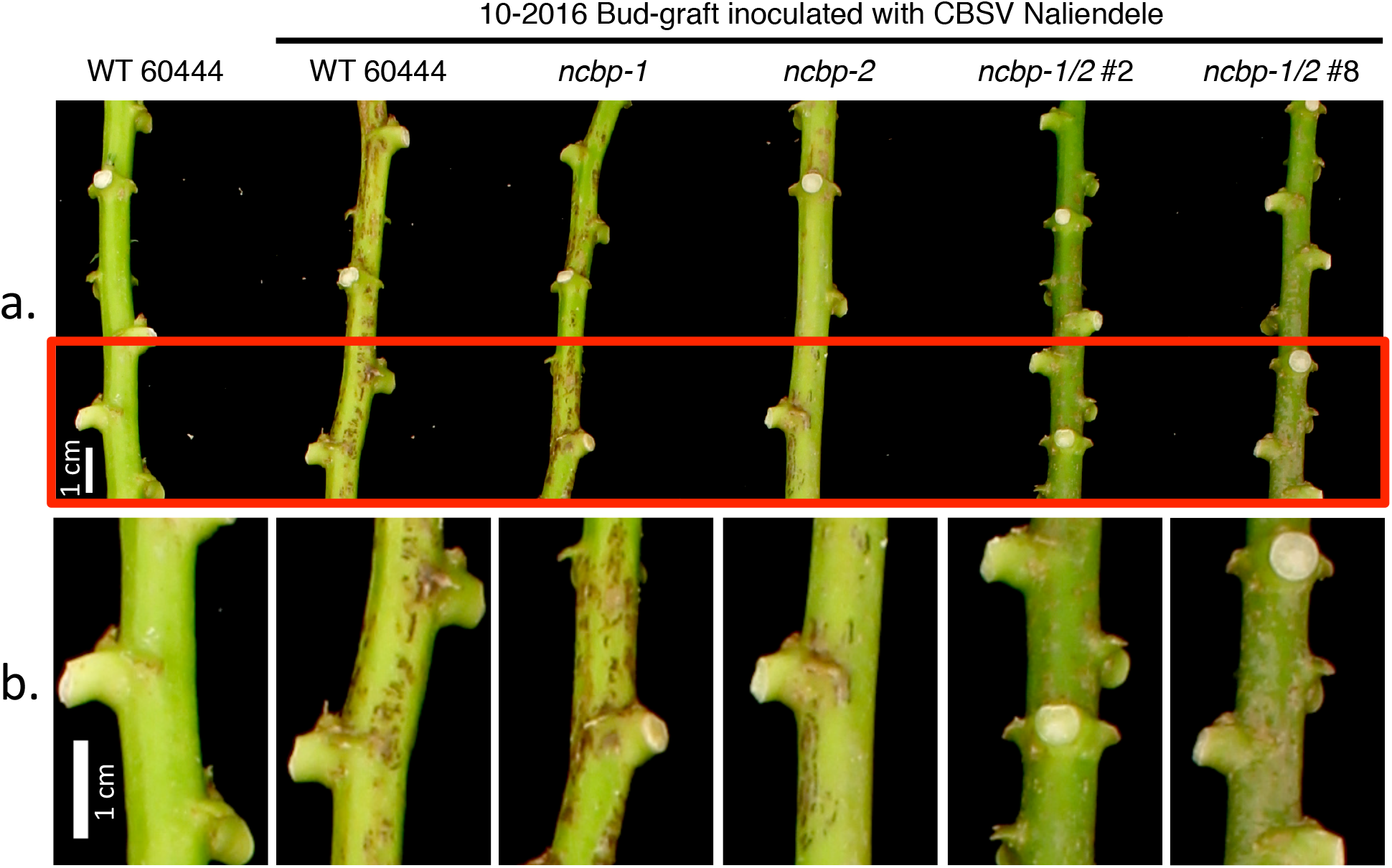
CBSD stem symptom attenuation on *ncbp-1 ncbp-2* double mutants. (a), representative wild type, *ncbp-1, ncbp-2*, or *ncbp-1 ncbp-2* stems displaying varying degrees of brown streak 14 weeks post graft inoculation with CBSV Naliendele. *ncbp-1 ncbp-2* double mutants present reduced brown streaking and associated dark pigmentation along the length their stems. Portions of stems boxed in red are enlarged in (b). Imaged portions of stems are all approximately the same distance from the graft site.

### *ncbp-1/ncbp-2* double mutant storage roots are less symptomatic and accumulate less virus

At 12 to 14 weeks after graft inoculation, storage roots were excavated and assessed for root necrosis. Each storage root of a plant was divided into five sections and each section scored on a 1-5 scale for CBSD symptom severity (Fig. 7a). Average symptom scores for each genotype were compared. *ncbp-2* and *ncbp-1/ncbp-2* mutant lines exhibited significantly reduced symptom scores relative to wild type and *ncbp-1* (Fig. 7a). Reverse transcription-<u>q</u>uantitative <u>p</u>olymerase <u>c</u>hain <u>r</u>eaction (qPCR) was used to measure CBSV-Nal RNA levels in *ncbp-1/ncbp-2* double mutants. Mean viral RNA levels in *ncbp-1/ncbp-2* roots were reduced 43-45% compared to wild-type roots (Fig. 7b).

**Figure 7.**
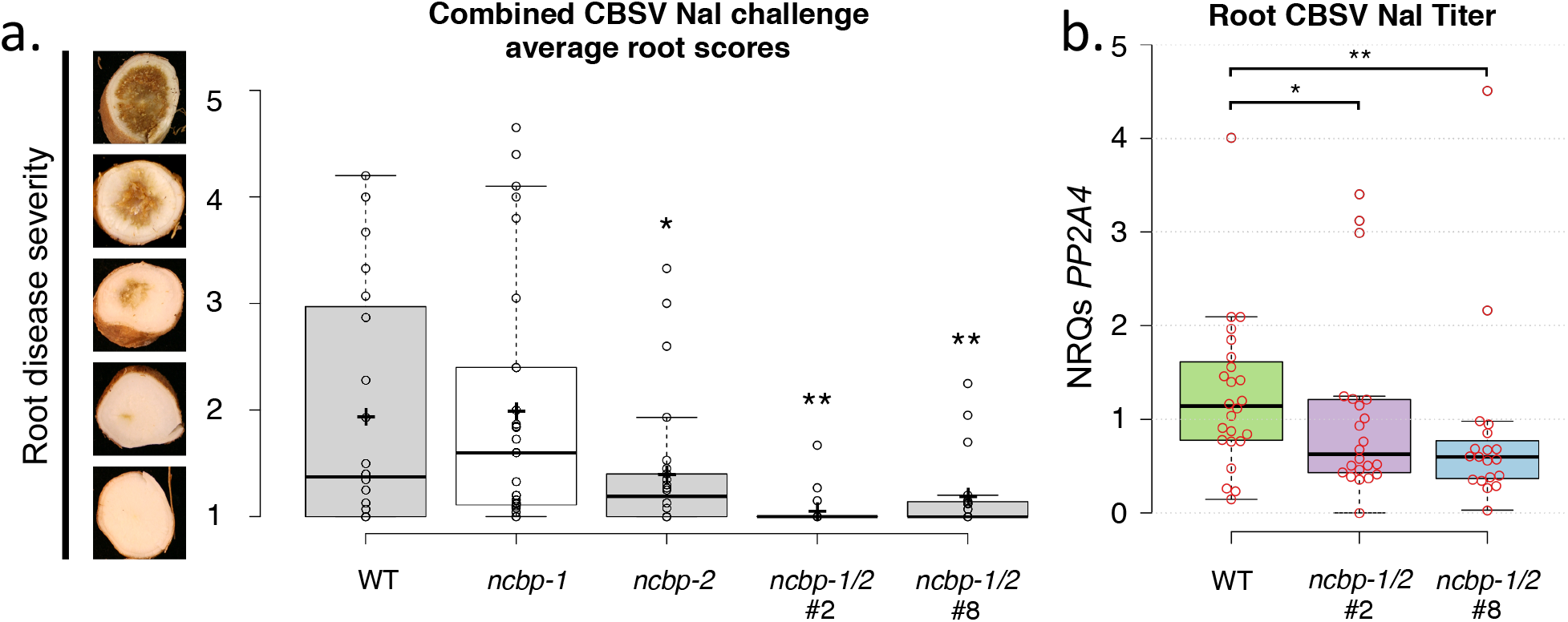
*ncbp-1 ncbp-2* double mutant storage roots are less symptomatic and accumulate less virus. (a), storage root sections were assessed on a 1-5 scale, 1 corresponding with asymptomatic and 5 corresponding with necrosis spanning the diameter of root. (b), ncbp-1 ncbp-2 storage roots are significantly less symptomatic than wild type at 12 to 14 weeks post bud-graft inoculation with CBSV Naliendele isolate. Points represent average scores of all storage root sections from a single plant. Data from three experimental replicates were pooled. Whiskers span the interquartile range, solid bars indicate the median of scores, + indicates the mean of scores. Statistical significance was detected by Welch’s t-test, n≥19, α=0.05, *≤0.05, **≤0.01, ****≤0.0001. (c), quantitative real time PCR analysis reveals that ncbp-1/2 storage roots accumulate less virus than wild type. CBSV *HAM1-LIKE* levels are reported as normalized relative quantities (NRQs) relative to *PP2A4.* Data from three experimental replicates were pooled. Significant differences were detected with a Mann-Whitney U-test, n≥19, α=0.05.

## Discussion

The CRISPR/Cas9 system has emerged as a powerful tool for plant genome editing and rapid crop improvement. In the context of disease resistance in crop species, this system has been employed to target *<u>mi</u>ldew-resistance <u>l</u>ocus <u>O</u> (MLO)* in wheat, and generate broad potyvirus resistance in cucumber by disrupting function of the *eIF4E* gene (Chandrasekaran *et al.*, 2016; Wang *et al.*, 2014). In the present study, we targeted the *nCBPs* to assess their putative function as CBSD susceptibility factors in cassava.

Previous studies have shown that host eIF4E and viral VPg interaction is necessary for potyviral infection (Ashby *et al.*, 2011; Charron *et al.*, 2008; Kang *et al.*, 2005; Leonard *et al.*, 2000; Yeam *et al.*, 2007). We identified five eIF4E family members in cassava, corroborating a recent analysis by Shi *et al.* (2017). Attempts to identify markers associated with CBSD resistance indicate that multiple loci are involved (Maruthi *et al.*, 2014; Masumba *et al.*, 2017). Examination of CBSD-resistant, -tolerant, and -susceptible cultivars by Shi and colleagues also found that these categories are not associated with *eIF4E* family single nucleotide polymorphisms (Shi *et al.*, 2017). As such, a biochemical study of the VPg and eIF4E family interaction was essential to identify a potential susceptibility gene(s).

Yeast two-hybrid and co-IP analysis showed consistent interactions between the nCBPs and the CBSV-Naliendele VPg (Fig. 2). nCBPs form a distinct clade from other eIF4E isoforms and demonstrate methylated-cap-binding property (Kropiwnicka *et al.*, 2015; Ruud *et al.*, 1998). There is no precedent for recruitment of nCBPs by potyviral VPg proteins. However, nCBP has been identified as a novel recessive resistance gene toward viruses in the *Alphaflexiviridae* and *Betaflexiviridae* families, specifically by inhibiting accumulation of movement proteins from a viral subgenomic RNA (Keima *et al.*, 2017). nCBP may be similarly involved in the accumulation of the CBSV movement protein, although differences in viral replication strategies suggest a different underlying mechanism (Hagiwara-Komoda *et al.*, 2016; Olspert *et al.*, 2015; Rodamilans *et al.*, 2015). Distantly related potyviruses with common hosts may also utilize different eIF4E isoforms for movement (Contreras-Paredes *et al.*, 2013; Eskelin *et al.*, 2011; Gao *et al.*, 2004; Miras *et al.*, 2017). Evidence also suggests that some potyviruses utilize one specific isoform for translation and another distinct isoform for movement (Contreras-Paredes *et al.*, 2013; Gao *et al.*, 2004). This complexity makes it difficult to predict what roles cassava nCBPs may have in the CBSV life cycle.

In our CRISPR/Cas9 edited lines, we observed homozygous, bi-allelic, heterozygous, complex, and wild-type genotypes. Homozygous mutations may have been generated by identical NHEJ repair, or homologous recombination-based repair from the opposite allele. Considering the low incidence of the latter in plants, identical NHEJ repair may be more likely (Peng *et al.*, 2016). While transgenic plants derived from FECs are thought to be of single cell origin (Fig. S3), reducing the likelihood of transgenic chimeras (Schreuder *et al.*, 2001; Taylor *et al.*, 1996), Odipo et al. (2017) have reported the production of chimeric plants via CRISPR/Cas9-mediated gene editing of phytoene desaturase in cassava. Complex genotypes were not characterized in depth, but they are likely chimeras resulting from Cas9/sgRNA activity being delayed until after embryogenic units began to replicate (Odipio *et al.*, 2017; Zhang *et al.*, 2014). Integrating CRISPR/Cas9 constructs into the cassava genome proved to be efficient for achieving gene editing as 91% of transformed lines carried INDELs at the target sites, and desired homozygous and bi-allelic mutations were observed in 78% of plant lines. These frequencies compare favorably to previous CRISPR/Cas9-mediated mutagenesis studies in cassava, rice and tomato (Ma *et al.*, 2015; Odipio *et al.*, 2017; Pan *et al.*, 2016).

A set of off-targets with five or fewer mismatches were also analyzed. A single mutation for one allele of off-target G was observed. Notably, this off target had the least number of mismatches: two consecutive nucleotides located furthest from the PAM. This type of mismatch has been demonstrated to be tolerated, but any additional mismatch within the protospacer eliminates off-target editing (Anderson et al., 2015). Congruent with this, the remaining wild-type allele of off-target G has an additional third mismatch. No further edits were detected for the remaining off-targets, with the exception of unsequenced off-target D which lies in an intergenic dinucleotide-rich locus. These data strongly suggest that our mutant phenotypes are due to on-target effects.

CBSV-Naliendele challenge experiments were performed on single-*ncbp* and double-*ncbp* mutant lines carrying homozygous and bi-allelic mutations that ultimately resulted in frameshifted coding sequences. Double-*ncbp* mutant lines exhibited delayed CBSD aerial symptom onset and reduced disease severity. Aerial virus titer was then quantified in endpoint leaf samples, but no significant differences between wild-type and double-*ncbp* mutants were detected. While stem symptom severity was significantly reduced in double-*ncbp* mutants throughout the entirety of each challenge, endpoint leaf symptom severity was comparable to wild-type in all challenges (Fig. S8). This may explain the lack of difference in leaf virus titer. Alternatively, it is well documented that virus accumulation in different aboveground organs can be uneven, regardless of uniformity in tissue type (Ogwok *et al.*, 2015). This may explain the large variances in virus titer observed within genotypes and impede detection of subtle yet significant differences. In contrast to leaf tissues, double-*ncbp* mutant storage roots exhibited significantly reduced levels of necrosis at the end of our challenges, comparable to observed differences in stem symptoms. This was also reflected by significantly reduced virus titers in storage roots of double-*ncbp* mutants. If future strategies result in a stronger level of resistance, we expect leaf virus titer to be more informative.

Single-*ncbp* mutants were not consistently significantly different from the susceptible wild-type plants in response to CBSV-Nal challenges. However, in several assays the *ncbp-2* mutant line showed an intermediate phenotype (Fig. S6a, S7a, S7c, S8a, S8b). In this work, three independent disease trials were performed in February, October and December of 2016. Although plants were maintained in greenhouse facilities, it is impossible to maintain completely consistent environmental conditions. These variations may account for the variability we observed across experiments. For example, in January 2017, wild-type stem severity scores peaked and were similar for challenges started at the end of 2016. Temperature fluctuations are also thought to influence CBSD leaf symptom presentation, with high temperatures inhibiting development of chlorosis (Hillocks and Jennings, 2003). These variations in disease pressure may have obscured the intermediate aerial phenotype of nCBP-2 mutants. In storage roots, however, mutagenesis of *nCBP-2* resulted in reduced symptom severity as compared to wild-type plants and *ncbp-1* mutant lines (Fig. 7). Notably, *nCBP-2* is expressed 10 fold more than *nCBP-1* in the storage roots, and remaining *4E* isoforms at levels less than *nCBP-1* (Fig. S11) (Wilson *et al.*, 2017). Assuming that *nCBP* mutations disrupt VPg-nCBP interactions, it is also possible that CBSV VPg may have a greater dependence for nCBP-2 than nCBP-1. Forcing CBSV to utilize less abundant isoforms, or those with suboptimal binding affinities, could attenuate CBSD progression (Chandrasekan et al., 2016). The high levels of *nCBP-2* in storage roots may also be a major contributing factor to development of root necrosis. Additional transcriptional and biochemical studies will be needed to investigate these hypotheses.

Several explanations may account for the incomplete CBSD resistance of double*-nCBP* mutant cassava plants. First, unpredicted splice variants may have coded for proteins that were biologically functional for viral infection, at least in part (Fig. 4). Disruption of the wild-type splice site motifs, typically dinucleotides GU and AG at the 5’ and 3’ termini of introns, respectively, can activate cryptic splice sites that redefine intron boundaries and consequently frameshift the mature transcript (Lal *et al.*, 1999, Reddy et al., 2007). This is consistent with our cDNA clone-seq analysis identifying a type 3 splice variant of *ncbp-1* from line *ncbp-1/2* #8. However, the type 2 *ncbp-1* variant of *ncbp-1/2* #2 does not appear to be the result of splice site disruptions and codes for full length nCBP-1 with a 12 amino acid extension. It is possible that similar unpredicted splice variants exist at low abundance in *ncbp-1/2* #8. Complementation assays will need to be performed to determine whether such putative splice variants can be utilized by the viruses. Experiments with edited lines that avoid splice sites can be generated to test this hypothesis. Alternatively, CBSV VPgs may have some affinity for the other eIF4E isoforms. Coexpression of the cassava eIF(iso)4E-1 and -2 with VPg from both species in yeast showed weak reporter activation that could be interpreted as weak interaction (Fig. 2). Co-IP analysis found that all cassava eIF4E isoforms could associate with CBSV-Nal VPg. VPg is an intrinsically disordered protein, which could enable it to bind several different proteins (Jiang and Laliberté, 2011). The ability to use multiple eIF4E isoforms has precedence, such as in *Pepper veinal mottle potyvirus* for which simultaneous mutations of both eIF4E and eIF(iso)4E are required to restrict infection (Gauffier *et al.*, 2016; Ruffel *et al.*, 2006). Further investigation will be required to determine the extent to which these other eIF4E isoforms contribute to CBSD.

CBSD remains a major threat to food security in sub-Saharan Africa. Mitigation of crop losses is imperative to sustaining Africa’s rapidly growing population. Due to the challenges of breeding cassava, gene editing strategies provide an attractive means to engineer disease resistance. Gene editing reagents integrated into the genome can be segregated away within an F1 population or possibly removed using site specific recombinase technology such as the Cre-lox system. In this study, we show that simultaneous CRISPR/Cas9-mediated editing of the *nCBP-1* and *nCBP-2* genes confers statistically significantly elevated resistance to CBSD. Editing of these host translation factors significantly hampers CBSV accumulation in the plant. By stacking this approach with other forms of resistance such as RNAi, potential exists to provide improved cassava varieties with robust and durable resistance to CBSD.

### Experimental Procedures

#### Production of plants and growth conditions

Transgenic cassava lines of cultivar 60444 were generated and maintained *in vitro* as described previously (Taylor *et al.*, 2012). *In vitro* plantlets were propagated, established in soil, and transferred to the greenhouse (Taylor *et al.*, 2012; Wagaba *et al.*, 2013). Throughout the course of a disease trial, all plants were treated bi-weekly for pest control by gently spraying the undersides of all leaves with water.

#### Identification and phylogenetic analysis of eIF4E isoforms

BLAST search of the AM560-2 cassava cultivar genome was done via Phytozome V10 using *A. thaliana* eIF4E family proteins as the queries. The coding sequences of each isoform were verified by comparison to RNA-seq data (Cohn *et al.*, 2014). Clustal Omega (EMBL-EBI) was used to generate the percent identity matrix of all eIF4E isoform amino acid sequences (Goujon *et al.*, 2010; Sievers *et al.*, 2014). MEGA 6 software was used to generate a phylogenetic tree of the cassava and *A. thaliana* eIF4E isoforms (Tamura *et al.*, 2013). The evolutionary history was inferred by using the Maximum Likelihood method based on the Le_Gascuel_2008 model (Le and Gascuel, 1993). This amino acid substitution model was determined as best fit using the MEGA 6 model test. The tree with the highest log likelihood (-2025.7966) is shown. Initial tree(s) for the heuristic search were obtained automatically by applying Neighbor-Join and BioNJ algorithms to a matrix of pairwise distances estimated using a JTT model, and then selecting the topology with superior log likelihood value. A discrete Gamma distribution was used to model evolutionary rate differences among sites (5 categories (+*G*, parameter = 1.9218)). The analysis involved 9 amino acid sequences. All positions containing gaps and missing data were eliminated.

#### CRISPR/Cas9 binary construct design

CRISPR/Cas9 construct design and assembly of entry clone pCR3 were conducted as described by Paula de Toledo Thomazella *et al.* (2016). CRISPR/Cas9 constructs targeting two sites were assembled via Gibson Assembly of the other U6-26/gRNA into the SacII site of the entry clone. Flanked by the attL1 and attL2 recombination sequences, the cassette carrying the Cas9/gRNA expression system was Gateway cloned into the binary destination vector pCAMBIA2300 (Hajdukiewicz *et al.*, 1994).

#### gRNA design and cloning

Target sequences were identified in *nCBP-1* and *nCBP-2* genes of cassava using the online CRISPR-P software (Lei *et al.*, 2014). This tool was used to select targets with predicted cut sites within exons, minimal off-target potential, and overlapping restriction enzyme recognition sites.

gRNA forward and reverse primers were designed with overhangs compatible with the BsaI-site described above. The Golden Gate (GG) cloning method was used to BsaI digest the pCR3 vector and ligate in the gRNA. In the case of the dual targeting CRISPR/Cas9 construct, the pCR3 vector bearing gRNA1 was digested with SacII, a site within the LR clonase attL sequences. The A. *thaliana* U6-26 promoter and gRNA2 were PCR amplified using primers suitable for Gibson Assembly into the SacII cut site of the digested pCR3-gRNA1 vector. For Gibson Assembly, 100 ng of SacII-digested vector was incubated with 200 ng of U6-26p-gRNA2 PCR amplicon and Gibson Assembly Master Mix for one hour and transformed into *E. coli* (NEB5a). Sequences of cloned CRISPR constructs were verified via Sanger sequencing.

#### Yeast two-hybrid

The *eIF4E* isoforms were amplified by PCR using primers suitable for Gibson Assembly into the *BamHI* site of pEG202. Yeast codon optimized coding sequences of the CBSV and UCBSV VPg were synthesized through Genewiz, Inc (South Plainfield, NJ, USA). The VPg coding sequences were amplified using primers suitable for Gibson Assembly into the *EcoRI* site of pEG201. Yeast two-hybrid analyses were carried out as described previously (Kim *et al.*, 2014).

#### Recombinant CBSV VPg purification

To generate a CBSV Naliendele VPg-3xFLAG fusion, CBSV VPg was cloned from cDNA into pDONR207 (ThermoFisher) and recombined into pK7-HFC (Huang et al., 2015). *CBSV VPg-6xHis-3xFLAG* was then cloned from this vector into pENTR/D-TOPO (Life) and recombined into the pDEST17 expression vector. Protein expression and purification was performed as described in Lin et al., 2015 with the following modifications. pDEST17-CBSV VPg-6xHis-3xFLAG was transformed into *Escherichia coli* BL21-AI cells. Protein expression in two liters of culture, optical density at 600 nm = 0.4, was induced with arabinose (0.2%) at room temperature for four hours. Clarified lysate was supplemented 100 units of DNAse I (ThermoFisher) and briefly sonicated to reduce viscosity. After incubation of Ni-NTA beads with cell lysate, beads were washed as previously described and CBSV VPg was eluted with four stepwise elutions of 100 mM imidazole followed by three stepwise elutions of 500 mM imidazole.

#### Co-immunoprecipitation

Cassava *eIF4E* isoforms were cloned from 60444 cDNA into pENTR/D-TOPO and recombined into pEARLEYGATE104. A *35S::YFP* control was made by removing the pEARLEYGATE104 ccdB cassette with BamHI and XbaI. Resulting overhangs were blunted and ligated to generate pEARLEYGATE104 ccdB-. These constructs were transformed into *Agrobacterium tumefaciens* strain GV3101. YFP and YFP-fusion proteins were co-expressed with p19-HA (Chapman et al., 2004) in *Nicotiana benthamiana* leaves. 48 hours post infiltration, leaves were harvested, frozen in liquid nitrogen, ground to fine powder, and resuspended with 2 mL of immunoprecipitation buffer (50 mL Tris-Cl pH=7.5, 150 mM NaCl, 10 mM MgCl2, 0.1% IGEPAL, 1x cOmplete EDTA-free protease inhibitor, 10 mM DTT) per gram of tissue. Leaf extract was centrifuged at 4000g for 15 minutes to remove debris. 1.2 mL of supernatant was then incubated with 6 ug 6xHIS-CBSV VPg-6xHIS-3xFLAG and 15 uL GFP-Trap magnetic agarose beads (ChromoTek) for 1.5 hours. Beads were washed six times with 1 mL immunoprecipitation buffer and boiled in 45 uL Laemmli sample buffer to elute bound proteins. Input and eluate were analyzed by immunoblotting with anti-GFP (Abcam ab290) and anti-FLAG M2 peroxidase (Sigma).

#### Genotyping and mutant verification

100 mg of leaf tissue was collected from T0 transgenic cassava *in vitro* plantlets and genomic DNA extracted using the CTAB extraction procedure (Murray and Thompson, 1980). Transgenic plants were genotyped for Cas9-induced mutagenesis via <u>r</u>estriction <u>e</u>nzyme <u>s</u>ite <u>l</u>oss (RESL) and Sanger sequencing (Voytas, 2013). Initially, 100 ng of genomic DNA was PCR amplified using primers encompassing the Cas9 target sites. PCR amplicons were gel purified on 1.5% agarose gel and purified with the QIAquick Gel Extraction Kit. For RESL analysis, 50 ng of PCR amplicon were digested with restriction enzyme SmlI for 12 hours, then run and visualized on a 1.5% agarose gel. For genomic and cDNA sequence analysis, the amplicons were subcloned and Sanger sequenced through the UC Berkeley DNA Sequencing Facility. Between six to eight clones were sequenced to discriminate INDEL polymorphisms and sequences were aligned to the intact *nCBP-1* and *nCBP-2* using SnapGene software (from GSL Biotech; available at snapgene.com).

#### Off-target analysis

The software CasOT with default settings was used to search potential off target sites of the CRISPR/Cas9 system against the 60444 illumina sequenced genome mapped onto the AM560-2 reference genome (Bredeson et al., 2016). This reference-based assembly was created using CLC Genomics Workbench with the following parameters (match score = 3; mismatch cost = 1; insertion cost = 1; deletion cost = 1; length fraction = 0.8; Similarity fraction = 0.8; global alightment = yes). 318,428,498 reads were successfully mapped. Off-target sequences were examined by Sanger sequencing of PCR amplicons encompassing the off-targets.

#### CBSV inoculation and disease scoring

Prior to virus challenge, micropropagated cassava plantlets were transplanted to soil, allowed to acclimate for six to eight weeks, and chip-bud graft inoculation performed as described previously (Beyene *et al.*, 2017; Wagaba *et al.*, 2013). Briefly, one plant of each genotype received an axillary bud from a single previously infected wild type plant, resulting in one inoculation cohort. Multiple cohorts were used in a single experiment to control for donor plants with varying viral concentrations.

Shoot tissues were scored two to three times a week for 12 to 14 weeks. Leaves and stems were each scored on separate 0-4 scales (Table S2). Average of leaf and stem scores are displayed for each time point. These data were used to calculate the area under the disease progression curve (Simko and Piepho, 2012). To assess symptom severity in storage roots, each storage root was evenly divided into five pieces along its length. Each storage root piece was then sectioned into one-centimeter slices and the maximum observed severity was used to assign a symptom severity score to that storage root piece. The scores for all storage root pieces of a given plant were then averaged to determine the overall severity score.

#### Storage root viral titer quantification

Five to ten representative storage root slices per plant were collected for viral titer quantification. Samples were flash frozen in liquid nitrogen and lyophilized for two days. Lyophilized storage roots were pulverized in 50 mL conical tubes with a FastPrepTM-24 instrument (MP Biomedicals) and 75 mg of pulverized tissue was aliquoted into Safe-Lock microcentrifuge tubes (Eppendorf) pre-loaded with two mm zirconia beads (BioSpec Products). Samples were flash frozen in liquid nitrogen, and further homogenized to a finer consistency. One mL of Fruit-mate (Takara) added to each sample. Samples were homogenized and subsequently centrifuged to remove debris. The supernatant was removed, mixed with an equal volume of TRIzol LS (Thermo Fisher), and the resulting mixture processed with the Direct-zol RNA MiniPrep kit (Zymo Research). Resulting total RNA was normalized to a standard concentration and used for cDNA synthesis with SuperScript III reverse transcriptase (Thermo Fisher).

Quantitative PCR was done with SYBR Select Master Mix (Thermo Fisher) on a CFX384 Touch Real-Time PCR Detection System (Bio-Rad). Primers specific for CBSV-Nal *HAM1-LIKE* and cassava *PP2A4* (Manes.09G039900) were used for relative quantitation (Table S5). Normalized relative quantities were calculated using formulas described by Hellemans *et al.* (2007). For combined analysis of all experimental replicates, normalized relative quantities for all samples were further normalized as a ratio to the geomean of wild type for their respective experiments. Data were then pooled and a Mann-Whitney U test was used to assess statistical differences.

## Acknowledgments

We are grateful for valuable advice and discussion with members of the VIRCA group. We thank Claire Albin, Maxwell Braud, the Staskawicz lab staff, the DDPSC tissue culture facility, and the DDPSC greenhouse staff for their work supporting this study. Funding in the Staskawicz Laboratory was provided by the Two Blades Foundation. Funding to R.B. and J.C. from the Bill & Melinda Gates Foundation (OPP1125410). K.R. was funded by the DDPSC NSF-REU program (DBI-1659812). The authors declare no conflicts of interest.

## Legends

Table S1. Genotypes of all transgenic T0 cassava lines. WT, wild-type alleles; bi-allelic, two different mutated alleles; heterozygous, wild-type and mutated alleles; chimeric, more than two mutated alleles. d# and i# refer to deletions and insertions, respectively, with the number of bases mutated denoted by #. Highlighted transgenic events were used in CBSV/UCBSV challenge assays.

Table S2. Potential off-targets for gRNA1 and gRNA2.

Table S3. Analysis of gRNA1 and gRNA2 off-targets.

Table S4. Aerial symptom scoring scale.

Table S5. Primers used in this study.

Figure S1. Roles of host eIF4E-potyvirus VPg interaction and sources of recessive resistance. (a) Linkage of potyvirus VPg to its binding site on eIF4E can provide translation initiation via recruitment of necessary factors and ribosomal subunits, genomic stability via protection from host-encoded exonucleases, and intracellular trafficking via eIF4G microtubule binding activity. (b) Non-conservative amino acid changes and gene deletions that abolish VPg-eIF4E binding removes above described roles, therefore conferring recessive resistance

Figure S2. TuMV VPg purified from *E. coli* can associate with plant eIF(iso)4E *in planta.* Immunoprecipitation of *YFP-Arabidopsis* eIF(iso)4E, but not YFP alone, co-immunoprecipitates TuMV VPg-3xFLAG. YFP and *YFP-Arabidopsis* eIF(iso)4E were co-expressed with *Tomato bushy stunt virus* p19 in *Nicotiana benthamiana* leaves and harvested 48 hours post agroinfiltration. 6xHIS-VPg-6xHIS-3xFLAG was purified from *E. coli* and 6 ug of protein was added to clarified leaf extracts.

Figure S3. Method for generating CRISPR/Cas9 mediated gene edited cassava. (a) Transgenic cassava are produced via Agrobacterium-mediated transformation of friable embryogenic calli (FEC). 1) FEC are induced from somatic tissues by placing the latter on growth media supplemented with picloram. FEC are comprised of aggregated spheroid embryogenic units. Individual units (boxed in panel 1 and enlarged in panel 2) range from a few cells to 1 mm in diameter. 2) FEC are transformed with CRISPR/Cas9 constructs through coculture with Agrobacterium tumefaciens. Red semi-circles denote TDNA fragments and red spheres denote transformed cells. 3) Cells on the surface of embryogenic units, transformed or untransformed, divide to produce new embryogenic units. CRISPR/Cas9 editing can occur prior to or after division. Edited cells are colored purple. 4) Antibiotic selection kills mother and untransformed daughter embryoids. Dead cells marked with “X”. Transformed embryogenic units are spread over selective media and form colonies. One mature embryo per colony is recovered (5), and develops into a plantlet (6). Each regenerated plant is clonally propagated and referred to as a mutant line. (b) Workflow for mutant genotype characterization and line selection.

Figure S4. CRISPR/Cas9-induced mutagenesis evident in *nCBP-1* (a) and *nCBP-2* (b) via restriction enzyme site loss (RESL). PCR amplicons of targeted regions were digested with SmlI. Map of amplicons with nCBP exon (purple), protospacer adjacent motif (red), gRNA spacer (green), predicted Cas9 cut site (black arrow), and overlapping SmlI restriction enzyme recognition site (bold, red). Cassava is diploid, carrying two copies of each nCBP gene. Absence of the wild-type digested product indicates that both alleles were successfully mutagenized. Bands are measured relative to O’Gene Ruler 1 kb Plus Ladder. Experimental banding pattern is consistent with predicted RESL.

Figure S5. CRISPR/Cas9–induced mutagenesis creates out of frame mRNAs. Exon 1 and exon 2 splice junction of *nCBP-1* (a) and *nCBP-2* (b) were examined via sequence analysis of cDNA. Frameshifting is observed for all *ncbp* single and double mutants. Predicted Cas9 cut site is shown as a black arrow. STOP codon is shown as starred, red box.

Figure S6. *ncbp-1 ncbp-2* double mutants exhibit slowed CBSV symptom onset. (a), (b), CBSV aerial symptom incidence for challenges initiated in February and December of 2016, respsectively. *ncbp* double mutants consistently exhibit delayed symptom onset relative to wild-type and single mutants. Incidence is reported as percent of wild type, *ncbp-1, ncbp-2*, or *ncbp-1 ncbp-2* plants, bud-graft inoculated with CBSV Naliendele (n≥7), displaying any level of leaf chlorosis or stem streaking.

Figure S7. *ncbp-1 ncbp-2* double mutants exhibit reduced aerial CBSV symptom severity. (a), (b), disease progression curves of wild type, ncbp-1, ncbp-2, or ncbp-1 ncbp-2 plants bud-graft inoculated with CBSV Naliendele (n≥7). Leaf and stem symptoms were each scored on a 0-4 scale and averaged to obtain an aerial score. (c), (d), average area under the disease progression curve (AUDPC) derived from data plotted in (a) and (b). Error bars indicate standard error of the mean. Statistical differences were detected by Welch’s t-test, α=0.05, *≤0.05, **≤0.01, ****≤0.0001.

Figure S8. *ncbp-1/ncbp-2* stem symptom severity is consistently reduced across all experiments. Separate leaf and stem disease progression curves for wild type, *ncbp-1, ncbp-2*, or *ncbp-1 ncbp-2* plants bud-graft inoculated with CBSV Naliendele (n≥5). Leaf and stem symptoms were each scored on a 0-4 scale. (a), (c), and (e) represent leaf disease progression curves from three different experiments while (b), (d), and (f) represent corresponding stem disease progression curves. Error bars represent standard error of the mean.

Figure S9. 12-2016 CBSV challenge leaf symptom severity is similar across all genotypes. Wild type, *ncbp-1, ncbp-2*, or *ncbp-1 ncbp-2* plants bud-graft inoculated with CBSV Naliendele isolate all develop widespread chlorotic leaf symptoms. Leaf images were taken near 12-2016 challenge endpoint. Scale bar denotes one centimeter.

FigureS10. Leaf CBSV Naliendele quantitation at challenge enpdoint does not reveal consistent differences in foliar virus titer. Quantitative real time PCR analysis of endpoint CBSV-Naliendele titer in wild type, *ncbp-1/2* #2, and *ncbp-1/2* #8 leaf tissue. Leaf samples were collected from the first fully expanded leaf. CBSV *HAM1-LIKE* was normalized to *PP2A4* and *GTPb (Manes.09G039900* and *Manes.09G086600).* n≥5 per genotype. Whiskers span the interquartile range, solid bars indicate the median of scores. Significant differences were detected with a Mann-Whitney U-test, n≥5, α=0.05, *<0.05.

Figure S11. *nCBP-2* is highly expressed in storage roots. Heat map describing tissue specific expression of cassava eIF4E isoforms. *nCBP-2* is expressed roughly 10 to 45 fold more than other *eIF4E* isoforms in storage roots. Data was extracted from the Bart Lab Cassava Atlas (http://shiny.danforthcenter.org/cassava_atlas/). Expression values are defined as fragments per kilobase of transcript per million mapped reads (FPKM).

